# The puzzle of metabolite exchange and identification of putative octotrico peptide repeat expression regulators in the nascent photosynthetic organelles of *Paulinella chromatophora*

**DOI:** 10.1101/2020.08.26.269498

**Authors:** Linda Oberleitner, Gereon Poschmann, Luis Macorano, Stephan Schott-Verdugo, Holger Gohlke, Kai Stühler, Eva C. M. Nowack

**Author notes:** **Correspondence to** Eva C. M. Nowack.

## Abstract

The cercozoan amoeba *Paulinella chromatophora* contains photosynthetic organelles - termed chromatophores - that evolved from a cyanobacterium, independently from plastids in plants and algae. Despite the more recent origin of the chromatophore, it shows tight integration into the host cell. It imports hundreds of nucleus-encoded proteins, and diverse metabolites are exchanged across the two chromatophore envelope membranes. However, the limited set of chromatophore-encoded transporters appears insufficient for supporting metabolic connectivity or protein import. Furthermore, chromatophore-localized biosynthetic pathways as well as multiprotein complexes include proteins of dual genetic origin, suggesting coordination of gene expression levels between chromatophore and nucleus. These findings imply that similar to the situation in mitochondria and plastids, nuclear factors evolved that control metabolite exchange and gene expression in the chromatophore. Here we show by mass spectrometric analyses of enriched insoluble protein fractions that, unexpectedly, nucleus-encoded transporters are not inserted into the chromatophore inner envelope membrane. Thus, despite the apparent maintenance of its barrier function, canonical metabolite transporters are missing in this membrane. Instead we identified several expanded groups of short chromatophore-targeted orphan proteins. Members of one of these groups are characterized by a single transmembrane helix, and others contain amphipathic helices. We hypothesize that these proteins are involved in modulating membrane permeability. Furthermore, we identified an expanded family of chromatophore-targeted helical repeat proteins. These proteins show similar domain architectures as known organelle-targeted octotrico peptide repeat expression regulators in algae and plants suggesting their convergent evolution as nuclear regulators of gene expression levels in the chromatophore.

**Importance:** The endosymbiotic acquisition of mitochondria and plastids >1 billion years ago was central for the evolution of eukaryotic life. However, owing to their ancient origin, these organelles provide only limited insights into the initial stages of organellogenesis. The chromatophore in *Paulinella* evolved ~100 million years ago and thus, offers the possibility to gain valuable insights into early stages and common rules in organelle evolution. Critical to organellogenesis appears to be the establishment of nuclear control over metabolite exchange and gene expression in the endosymbiont. Here we show that the mechanism generating metabolic connectivity of the chromatophore fundamentally differs from the one for mitochondria and plastids, but likely rather resembles the poorly understood mechanism in various bacterial endosymbionts in plants and insects. Furthermore, we describe a novel class of chromatophore-targeted helical repeat proteins which evolved convergently to plastid-targeted expression regulators and are likely involved in gene expression control in the chromatophore.

## Introduction

Endosymbiosis has been a major driver for the evolution of cellular complexity in eukaryotes. During organellogenesis, linkage of the previously independent biological networks of the former host and endosymbiont resulted in a homeostatic and synergistic association. Two critical factors during this dauntingly complex process appear to be the establishment of metabolic connectivity between the symbiotic partners, and nuclear control over protein levels within the organelle.

Besides mitochondria and primary plastids that evolved via endosymbiosis >1 billion years ago, recently, a third organelle of primary endosymbiotic origin has been identified (1, 2). The photosynthetically active ‘chromatophore’ of cercozoan amoebae of the genus *Paulinella* evolved around 100 million years ago from a cyanobacterium (3, 4). Hence, scrutiny of *Paulinella* can help to determine the degrees of freedom in the integration process of a eukaryotic organelle. Similar to the evolution of mitochondria and plastids, also in the chromatophore, reductive genome evolution resulted in the loss of many metabolic functions (5, 6), around 70 genes were transferred from the chromatophore to the nucleus (7–9), and functions lost from the chromatophore genome are compensated by import of nucleus-encoded proteins (10, 11). In a previous study, we identified by mass spectrometry (MS) around 200 nucleus-encoded, chromatophore-targeted proteins in *Paulinella chromatophora* that fall into two classes (10). Short import candidates [<90 amino acids (aa)] lack obvious targeting signals, whereas long import candidates (>250 aa) carry a conserved N-terminal sequence extension ─ likely a targeting signal ─ that is referred to as ‘chromatophore transit peptide’ (crTP). Bioinformatic identification of crTPs allowed to extend the catalogue of import candidates to >400 proteins (10).

Metabolic capacities of chromatophore and host cell are complementary resulting in the need for extensive exchange of metabolites such as sugars, amino acids, and cofactors across the two envelope membranes that surround the chromatophore (5, 10, 12). Furthermore, substrates for carbon, sulfur, and nitrogen assimilation (e.g. HCO_3_^−^, SO_4_^2−^, NH_4_^+^) and metal ions (e.g. Mg^2+^, Cu^2+^, Mn^2+^, and Co^2+^) that serve as cofactors of chromatophore-localized proteins have to be imported into the chromatophore. Whereas the chromatophore inner membrane (IM) clearly derives from the cyanobacterial plasma membrane, the outer membrane (OM) has been interpreted as being host-derived (13, 14). The nature of the transporters underlying the deduced solute (and protein) transport processes across this membrane system is unknown.

In plants and algae, transport across the plastid IM is mediated by a large set of multi-spanning transmembrane (TM) proteins that are highly specific for their substrates. These transporters contain usually four or more TM α-helices (TMHs) and are of the single subunit secondary active or channel type (15). This set of transporters apparently evolved mainly via the retargeting of existing host proteins to the plastid IM rather than the repurposing of endosymbiont proteins (15–17). Transport across the plastid OM is enabled largely by (semi-)selective pores formed by nucleus-encoded β-barrel proteins (18).

Another important issue during organellogenesis is the establishment of nuclear control over organellar gene expression supporting (i) adjustment of the organelle to the physiological state of the host cell, and (ii) assembly of organelle-localized protein complexes composed of subunits encoded in either the organellar or nuclear genome in stoichiometric amounts (19, 20). Also in *P. chromatophora*, the import of nucleus-encoded proteins resulted in protein complexes of dual genetic origin [e.g. photosystem I (11)]. The difference in copy numbers between chromatophore and nuclear genome (~100 vs one or two copies) (7) calls for coordination of gene expression between nucleus and chromatophore.

To test the hypotheses that nuclear factors were recruited to establish (i) metabolic connectivity between chromatophore and host cell and (ii) control over gene expression levels within the chromatophore, here we analyzed the previously obtained proteomic dataset derived from isolated chromatophores and a newly generated proteomic dataset derived from enriched insoluble chromatophore proteins with a focus on chromatophore-targeted TM proteins and putative expression regulators.

## Results

### Paucity of chromatophore-encoded solute transporters

Although diverse metabolites have to be exchanged constantly between the chromatophore and cytoplasm, we identified genes for only 25 solute transporters on the chromatophore genome (5, 10, 12). As judged from the localization of their cyanobacterial orthologs, only 19 of these transporters putatively localize to the chromatophore IM [**Fig. 1**, **Table S1**]. In comparison, in *Synechococcus* sp. WH5701, a free-living relative of the chromatophore (21), and the model cyanobacterium *Synechocystis* sp. PCC6803, genes for ~89 and >100 putative envelope transporters were identified, respectively [**Fig. 1**, **Table S1**, (22)]. Substrates of most of the chromatophore IM transporters are – according to annotation – restricted to inorganic ions (e.g. Na^+^, K^+^, Fe^2+^, Mg^2+^, PO_4_^2−^, HCO_3_^−^). Notably, cyanobacterial uptake systems for nitrogen and sulfur compounds such as nitrate (23), ammonium (24), urea (25), amino acids (26) or sulfate (27) are missing. Only one transporter of the DME-family (10 TMS Drug/Metabolite Exporter; PCC0734) could potentially be involved in metabolite export, and one transporter of the DASS-family (Divalent Anion:Na^+^ Symporter; PCC0664) could facilitate import of either di-/tricarboxylates or sulfate via Na^+^ symport. However, due to the multitude of substrates transported by members of both families (28, 29), precise substrate specificities cannot be predicted. Chromatophore-encoded β-barrel OM pores could not be identified.

**Figure 1:**
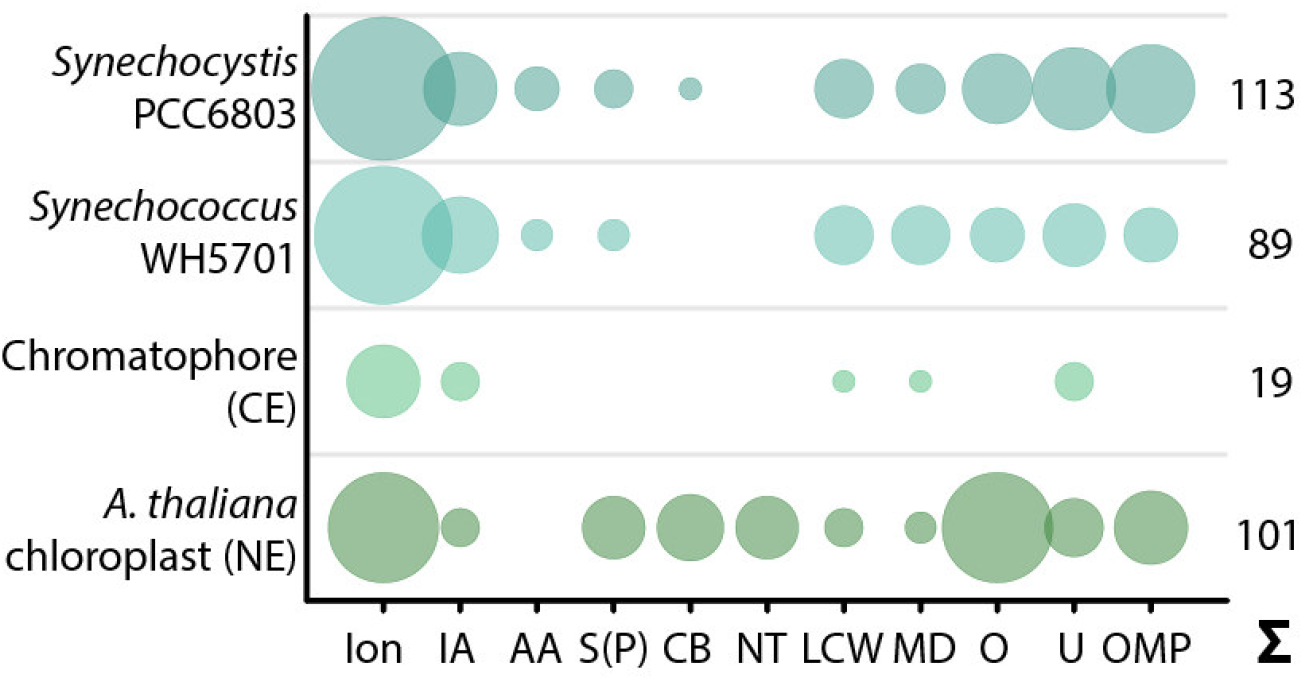
Predicted solute transport capacities of the chromatophore, *Synechococcus* sp. WH5701, *Synechocystis* sp. PCC6803, and the *Arabidopsis thaliana* chloroplast. Only transport systems for which experimental evidence suggests localization to the plasma membrane or the organellar envelope are shown. CE, chromatophore-encoded; NE, nucleus-encoded; Ion, ions/metals; IA, inorganic anions (phosphate, sulfate, nitrate, bicarbonate); AA, amino acid; S(P), sugars (hexoses, oligosaccharides) or sugar-phosphates; CB, mono-/di-/tricarboxylates; NT, nucleotides; LCW, lipid and lipopolysaccharide; MD, multidrug; O, other; U, unknown; OMP, outer membrane pores; Σ, total predicted transporters.

In contrast, in plants and algae, a combination of bioinformatic and proteomic studies identified 100-150 putative solute transporters in the plastid IM; 37 of these transporters have been confidently assigned functions and many of them transport metabolites [(16, 30–32), **Fig. 1**, and **Table S1**]. Several porins are known to permit passage of solutes across the chloroplast OM (18, 33–35). Almost all of these transport systems are encoded in the nucleus and posttranslationally inserted into the plastid envelope membranes.

### Enrichment of insoluble protein fractions and proteomic analysis

The scarcity of chromatophore-encoded solute transporters suggested that in *P. chromatophora*, as in plastids, nucleus-encoded transport systems establish metabolic connectivity of the chromatophore. However, among 432 previously identified import candidates (10), only 3 proteins contained more than one predicted TMH (**Table 1**). One of these proteins (identified by in silico prediction) contains two TMHs, only one of which is predicted with high confidence. Of the other two proteins (identified by MS), one is short and contains two predicted TMHs; the other contains eight predicted TMHs. However, this latter protein was identified with one peptide only and shows no BlastP hits against the NCBI nr database, whereas an alternative ORF (in the reverse complement) shows similarity to an NAD-dependent epimerase/dehydratase. Therefore, this latter protein likely represents a false positive (a false discovery rate of 1% was accepted in this analysis).

**Table 1:** No nucleus-encoded solute transporters among previously identified import candidates. The table lists numbers of proteins previously identified to be imported into the chromatophore [by in silico prediction (based on presence of a crTP), liquid chromatography coupled to tandem MS (LC-MS/MS), and total] (10) sorted by the number of predicted TMHs (outside of the crTP).

|  | 0 TMH | 1 TMH | >1 TMH |
| --- | --- | --- | --- |
| **In silico predicted (crTP)** | 289 | 3 | 1 |
| **LC-MS/MS identified** | 194 | 11 | 2 |
| **Total** | 416 | 13 | 3 |

The absence of multi-spanning TM proteins among import candidates could have two reasons. (i) These proteins might lack a crTP, impairing their prediction as import candidates. (ii) TM proteins are often underrepresented in LC-MS analyses owing to low abundance levels as well as unfavorable retention and ionization properties. In fact, our previous MS analysis identified 47% of the soluble but only 21% of TMH-containing chromatophore-encoded proteins (**Fig. 2C**).

**Figure 2:**
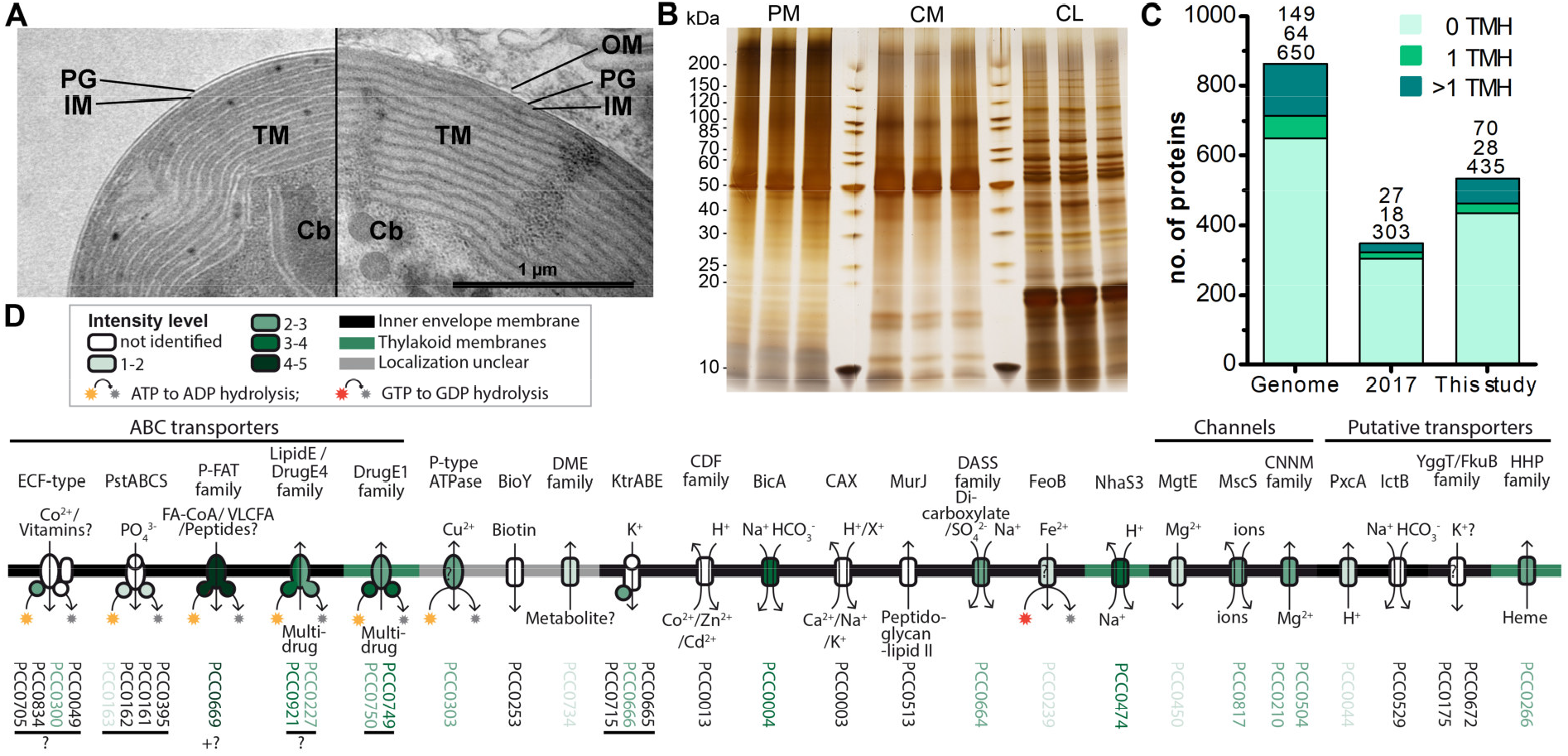
Increased recovery of TM proteins by MS analysis of enriched insoluble chromatophore proteins. **(A)** TEM micrographs of isolated chromatophore (left) and chromatophore in the context of a *P. chromatophora* cell (right). The outer envelope membrane (OM) observed in intact cells was lost during the isolation process. IM, inner envelope membrane; PG, peptidoglycan; TM, thylakoid membranes; Cb, carboxysomes. **(B)** 1 μg of protein from three replicates of each, chromatophore lysates (CL) as well as high salt and carbonate-washed *P. chromatophora* (PM) and chromatophore membranes (CM) was resolved on a 4-20 % polyacrylamide gel and silver stained. **(C)** Numbers of proteins encoded on the chromatophore genome (Genome) and chromatophore-encoded proteins identified with ≥3 SpC in chromatophore-derived samples in our previous (2017) and current (This study) proteome analysis. The number of predicted TMHs is indicated by a color code. **(D)** Detection of chromatophore-encoded transport systems. Annotation or TCDB-family, predicted mode of transport, substrates, and probable subcellular localization are provided. For each protein, the mean normalized intensity in CM (over both MS experiments) is indicated by a color code (see also **Table S1**).

Thus, to enhance identification of TM proteins, we enriched TM proteins by collecting the insoluble fractions from isolated chromatophores (CM samples) and intact *P. chromatophora* cells (PM samples). Electron microscopic analysis of isolated chromatophores suggested that the OM is lost during chromatophore isolation [**Fig. 2A**, compare (13, 14)]. Comparison of CM and PM samples to chromatophore lysates (CL samples) by SDS-PAGE revealed distinct banding patterns between the three samples and high reproducibility between three biological replicates (**Fig. 2B**). Further enrichment of membrane proteins or separation of IM, OM, and thylakoids was not feasible owing the slow growth of *P. chromatophora,* low yield of chromatophore isolations, and the loss of the OM. Two consecutive, independent MS analyses of three replicates of each, CM, PM, and CL samples led to the identification of 1,886 nucleus- and 555 chromatophore-encoded proteins over all fractions (**Table 2** and **S2**). Although most chromatophore-localized TM proteins were also identified in our analyses in CL samples (**Table 2**), individual TM proteins were clearly enriched in CM compared to CL samples (**Fig. S1**).

**Table 2:**
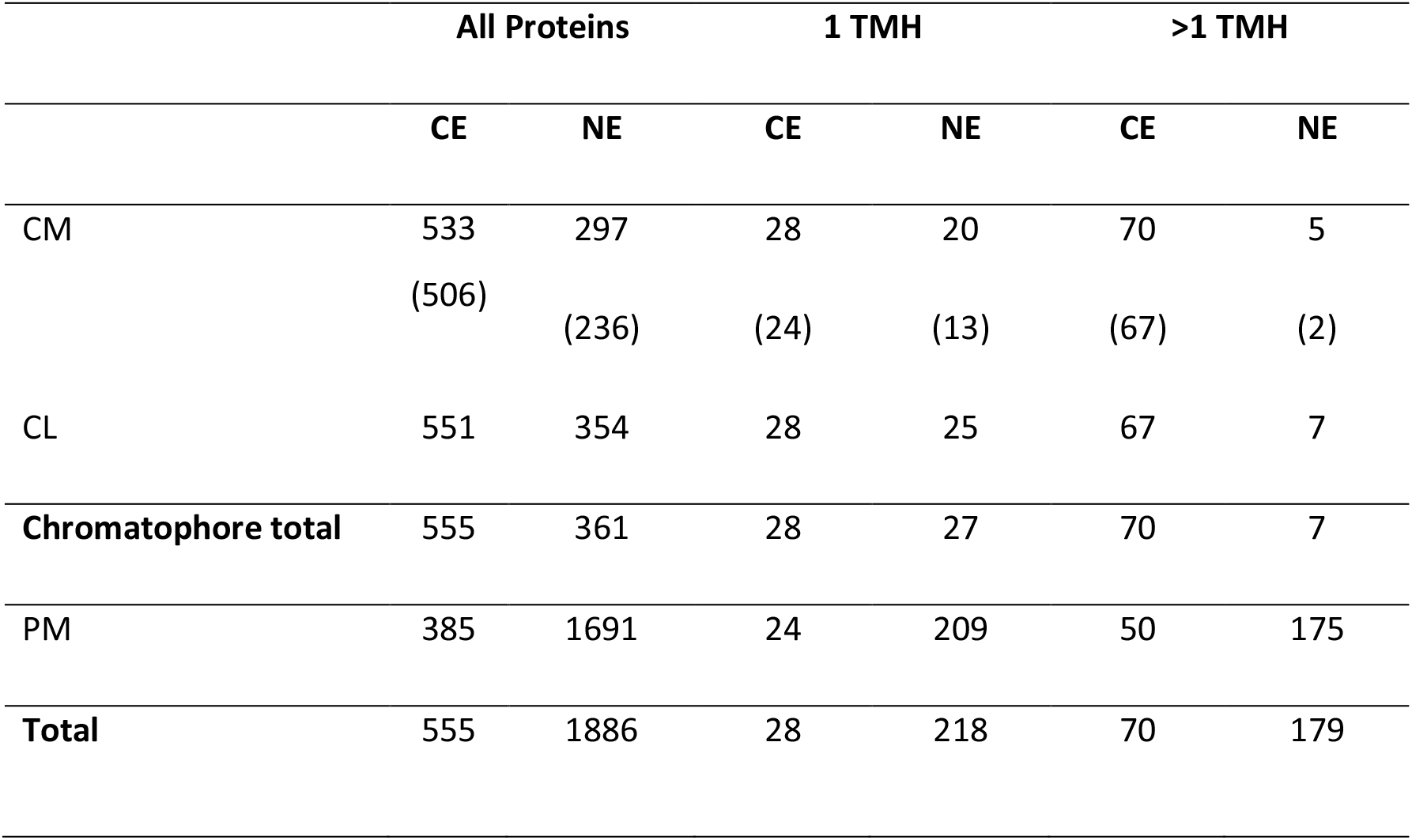
Proteins identified in this study by LC-MS/MS. Numbers of chromatophore-encoded (CE) and nucleus-encoded (NE) proteins identified in at least one out of two independent MS experiments with ≥3 spectral counts (SpC) in chromatophore-derived samples (i.e. CM+CL) or whole cell membranes (PM). The number of predicted TMHs (outside of the crTP) is indicated. For proteins identified in CM samples, total number of proteins and number of proteins enriched in CM as compared to PM samples (in brackets) is indicated separately.

|  | All Proteins |  | 1 TMH |  | >1 TMH |  |
| --- | --- | --- | --- | --- | --- | --- |
|  | CE | NE | CE | NE | CE | NE |
| CM | 533 | 297 | 28 | 20 | 70 | 5 |
|  | (506) | (236) | (24) | (13) | (67) | (2) |
| CL | 551 | 354 | 28 | 25 | 67 | 7 |
| **Chromatophore total** | 555 | 361 | 28 | 27 | 70 | 7 |
| PM | 385 | 1691 | 24 | 209 | 50 | 175 |
| **Total** | 555 | 1886 | 28 | 218 | 70 | 179 |

In CM samples, 46% (or 98/213) of the chromatophore-encoded TM proteins were identified, representing a gain of 118% compared to our previous analysis (**Fig. 2C**); in particular, of the 25 chromatophore-encoded solute transport systems, 72% (or 18) were identified with at least one subunit, and 60% (or 15) were identified with their TM subunit (**Fig. 2D**) while our previous study identified only three of these transporters. Highest intensities (representing a rough estimation for protein abundances) were found in CM samples for an ABC-transporter annotated as multidrug importer of the P-FAT family (level 4 to 5, placing the transporter among the 10% most abundant proteins in CM). Also the bicarbonate transporter BicA, two multidrug efflux ABC-transporters, and a NhaS3 proton/sodium antiporter were found in the upper tiers of abundance levels (level 3 to 4, placing them among the 30% most abundant proteins in CM). The remaining transporters showed moderate to low abundance levels (**Fig. 2D**).

### No multi-spanning TM proteins appear to be imported into the chromatophore

Determination of nucleus-encoded proteins enriched in CM compared to PM samples led to the identification of 188 high confidence (HC) [and further 48 low confidence (LC); see Methods and **Fig. S2**] import candidates (**Fig. 3A**, **Table S3**). Nucleus-encoded multi-spanning TM proteins appeared invariably depleted in chromatophores (**Figs. 3B**, **C**). Only two of 236 import candidates were multi-spanning TM proteins (**Table 2**). However, one of these (with 7 predicted TMHs, scaffold1608-m.20717, arrowhead in **Fig. 3B**) was identified by only one hepta-peptide and shows no similarity to other proteins in the NCBI nr database whereas an overlapping ORF (in another reading frame) encodes a peroxidase that was MS-identified in ref. (10) likely classifying the protein as a false positive. For the other import candidate (scaffold18898-m.107131; with an enrichment level close to 0; arrowhead in **Fig. 3C**) a full-length transcript sequence is missing precluding determination of the correct start codon. Thus, this protein might represent in fact a short import candidate with a single TMH. Of the three nucleus-encoded multi-spanning TM proteins that were present but appeared depleted in CM compared to PM samples (**Table 2**), two were annotated as mitochondrial NAD(P) transhydrogenase and mitochondrial ATP/ADP translocase, suggesting a mild contamination of CM samples with mitochondrial membrane material.

**Figure 3:**
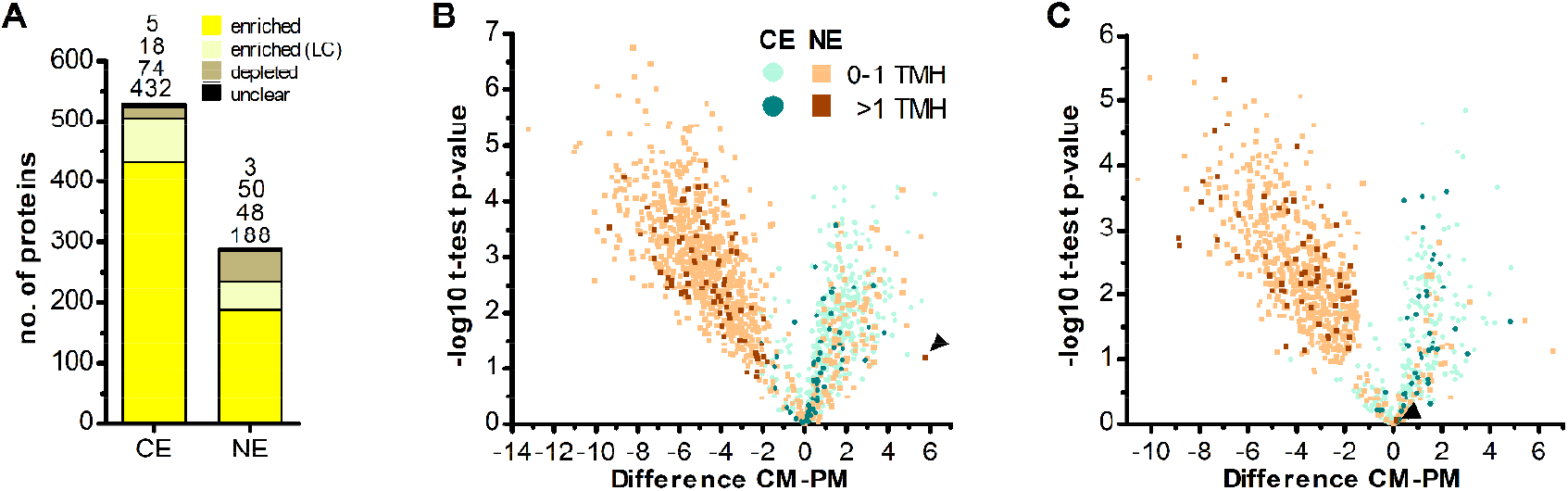
No evidence for import of host-encoded multi-spanning TM proteins into chromatophores. **(A)** Chromatophore-encoded (CE) and nucleus-encoded (NE) proteins enriched in CM compared to PM samples. Yellow, proteins enriched with high confidence; light yellow, proteins enriched with low confidence (LC); brown, proteins depleted in CM; black, proteins classified as “unclear” (see Methods and Fig. S2). Only proteins identified with ≥3 SpC in the chromatophore samples in at least one out of two independent MS experiments were considered. **(B, C)** The difference of intensities of individual proteins between CM and PM samples 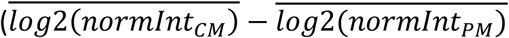; Difference) is plotted against significance (−log10 p-values in Student’s t-test) for proteins detected with ≥3 SpC in the chromatophore samples (for proteins detected in CM only or CM and PM) or in whole cell samples (proteins detected in PM only). Values for proteins detected only in one sample have been imputed and are only shown when their difference is significant. The number of predicted TMHs (outside of the crTP) is indicated by a color code. Data from MS experiment 1 **(B)** and 2 **(C)** are shown separately. Scaffold1608-m.20717 and scaffold18898-m.107131 (see text) are marked by arrowheads in B and C, respectively. In both analyses, among the proteins enriched in CM (Difference CM-PM > 0), the proportion of identified multi-spanning TM proteins encoded in the chromatophore (49 of 409 in B; 39 of 134 in C) as compared to the nucleus (0 of 132 excluding the false positive in B; 1 of 54 in C) is significantly higher (both: p-value = 0.002, Fishers’s Exact Test).

In comparison, 70 chromatophore-encoded multi-spanning TM proteins were identified in CM samples, and 67 of these appeared enriched in CM samples. In PM samples, 50 chromatophore- and 175 nucleus-encoded multi-spanning TM proteins were found (**Table 2**).

### Targeting of single-spanning TM proteins and antimicrobial peptide-like proteins to the chromatophore

In contrast to the striking lack of multi-spanning TM proteins, there were 13 (5 HC and 8 LC) single-spanning TM proteins (containing one TMH outside of the crTP) among the identified import candidates (**Table 2**). Three of these proteins contain a TMH close to their C-terminus and likely represent tail-anchored proteins. One of these proteins is long and annotated as low-density lipoprotein receptor-related protein 2-like, the other two (with N-terminal sequence information missing) as polyubiquitin. However, most import candidates with one TMH (10 proteins) represent short proteins. These short import candidates included two high light-inducible proteins [i.e. thylakoid-localized cyanobacterial proteins involved in light acclimation of the cell (9)]. The remaining eight proteins are orphan proteins lacking detectable homologs in other species (BlastP against NCBI nr database, cutoff e^−03^); all of these contain a TMH with a large percentage of small aa (26-45% Gly, Ala, Ser) close to their negatively charged N-terminus (**Fig. 4A**).

**Figure 4:**
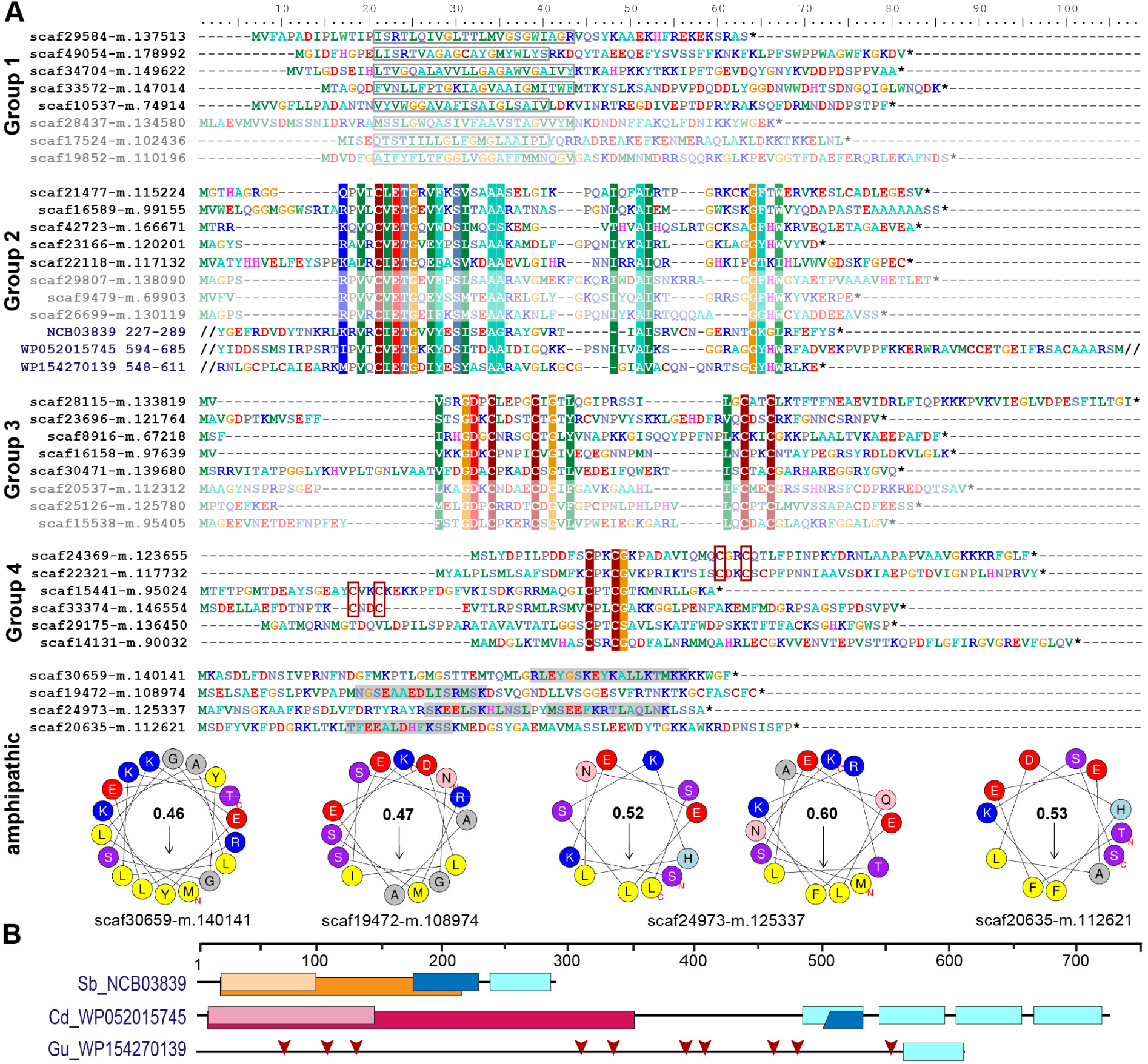
Short orphan import candidates form distinct groups. **(A)** For each group, representative MS-identified proteins (bright colors) and, if applicable, similar proteins identified among translated nuclear transcripts (pale colors) are displayed. Group 1: boxes indicate position of the predicted TMH. Group 2-4: colored background indicates ≥70% amino acid identity over alignments containing all MS-identified proteins of the respective group. The conserved sequence motif in group 2 was identified in diverse bacterial proteins (three examples with their NCBI accession number and aa positions are provided). Group 4: CxxC motifs are highlighted. Amphipathic: some short import candidates that do not belong to group 1 to 4 feature amphipathic helices. Areas highlighted in grey contain predicted alpha-helices. Corresponding wheel diagrams and hydrophobic moments are provided below. **(B)** Domain structure of bacterial proteins shown in A. Light blue boxes, conserved group 2 sequence motif; orange, group I intron endonuclease domain; light orange, GIY-YIG excision nuclease domain; pink, Superfamily II DNA or RNA helicase domain (SSL2); light pink, DEXH-box helicase domain of DEAD-like helicase restriction enzyme family proteins; blue, DNA-binding motif found in homing endonucleases and related proteins (NUMOD); red arrows, individual CxxC motifs. Sb, *Spirochaetia bacterium*; Cd, *Clostridioides difficile*; Gu, *Gordonibacter urolithinfaciens*.

In our previous proteome analysis, short orphan proteins represented the largest group of MS-identified import candidates (1/3 of total). However, most of these proteins did not possess predicted TMHs. Based on the occurrence of specific Cys motifs (CxxC, CxxxxC) and stretches of positively charged aas these short proteins were described as antimicrobial peptide (AMP)-like proteins (10). Including the eight TMH-containing proteins (see above), the recent study identified further 19 short orphan import candidates (or – only few proteins – showing similarity to hypothetical proteins in other species). Scrutiny of all 88 short orphan import candidates (resulting from both studies together) revealed that besides the TMH-containing proteins (group 1, 10 proteins), these short import candidates form at least three further distinct groups (**Fig. 4A**). Members of group 2 (12 proteins) contain a conserved motif of unknown function that occurs also in bacterial proteins that often possess domains pointing towards DNA processing-related functions (**Fig. 4A** and **B**). Members of group 3 (10 proteins) contain another conserved motif of unknown function that encompasses two Cys-motifs (CxxxxC and CxxC). Members of group 4 (30 proteins) show either one or two CxxC mini motifs (one of these is often CPxCG) but no further sequence conservation. The remaining 26 short orphan import candidates have no obvious common characteristics but several appear to have a propensity to form amphipathic helices (**Fig. 4A**).

Screening a large nuclear *P. chromatophora* transcriptome dataset (7) revealed additional putative members of groups 1 to 3 **(Fig. 4A** and **Fig. S3)**: further 53 translated transcripts represent short proteins with a predicted TMH in the N-terminal 2/3 of the sequence that is rich (>20%) in small aa and have an N-terminus with a net charge ≤0. Notably, the TMHs of >90% of all group 1 proteins comprise at least one (small)xxx(small) motif which can promote oligomerization of single-spanning TM proteins (36). Furthermore, many of these putative group 1 short import candidates are predicted to have antimicrobial activity and/or pore-lining residues (**Table S4**). Further 192 and 28 translated transcripts contain the conserved motifs of group 2 or 3, respectively. Importantly, all MS-identified members of these extended protein groups were identified in chromatophore-derived samples in this and our previous analysis.

### An expanded family of octotrico peptide repeat putative expression regulators is targeted to the chromatophore

Of the 235 import candidates (excluding the false positive, see above) identified in this study (**Fig. 3A**), 159 were known import candidates (10) (**Fig. 5A**, **Table S3**), with 46 proteins experimentally confirming import candidates previously only predicted in silico. 76 proteins represent new import candidates, mostly lacking N-terminal sequence information (42 proteins) or representing short import candidates (22 proteins). A particularly large number of newly MS-identified import candidates (24 proteins) fall into the category ‘genetic information processing’ (**Fig. 5B**). Among these proteins an expanded group of 10 RNA-binding or RAP domain-containing proteins [where RAP stands for **R**NA binding domain abundant in **ap**icomplexans, (37)] stood out.

**Figure 5:**
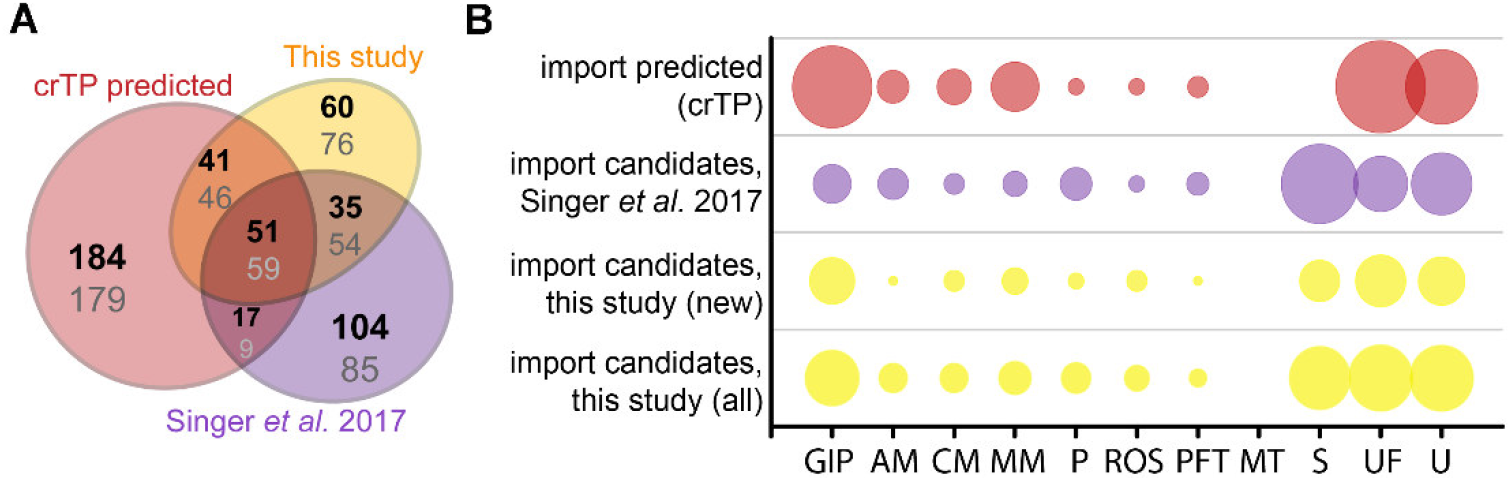
Newly identified import candidates. **(A)** Numbers of newly identified import candidates in this study [see Fig. 3A, yellow], previously MS-identified import candidates (Singer *et al.* 2017, purple), and in silico predicted import candidates [(10), red]. Numbers in bold indicate distribution of proteins considering only HC import candidates, numbers in grey considering all import candidates. **(B)** Functional categories of import candidates in (A). GIP, genetic information processing; AM, amino acid metabolism; CM, carbohydrate metabolism; MM, miscellaneous metabolism; P, photosynthesis and light protection; ROS, response to oxidative stress; PFT, protein folding and transport; MT, metabolite transport; S, short proteins (<90 aa) without functional annotation/homologs; UF, unspecific function; U, unknown function. “New” import candidates were MS-identified in this study, but not in Singer *et al.* 2017.

These RNA-binding proteins encompass, in addition to the crTP, from N- to C-terminus a variable region of 0-320 aa followed by a ~105 aa long conserved region (CR1), 2-13 repeats of a degenerate 38 aa motif with the most conserved residues being (xxxPxxxxLxxxxxxxxxxxxxFxxQxxxxxLNAxAKL), often followed by a 110 aa long conserved sequence (CR2), and the 60 aa long RAP domain (**Fig. 6**). This domain organization resembles the one of organelle-targeted octotrico peptide repeat (OPR; i.e. 38 aa peptide repeat) gene expression regulators in green algae and plants (**Fig. 6B, D**) and repeat-containing T3SS effector proteins described from symbiotic or pathogenic bacteria (**Fig. 6B, E, F**). All repeat motifs share the prediction to fold into two antiparallel α-helices. 3D-structure prediction suggests folding of the α-helical repeats in *Paulinella* OPR proteins into a super helix (or α-solenoid) structure (**Fig. 6G**).

**Figure 6:**
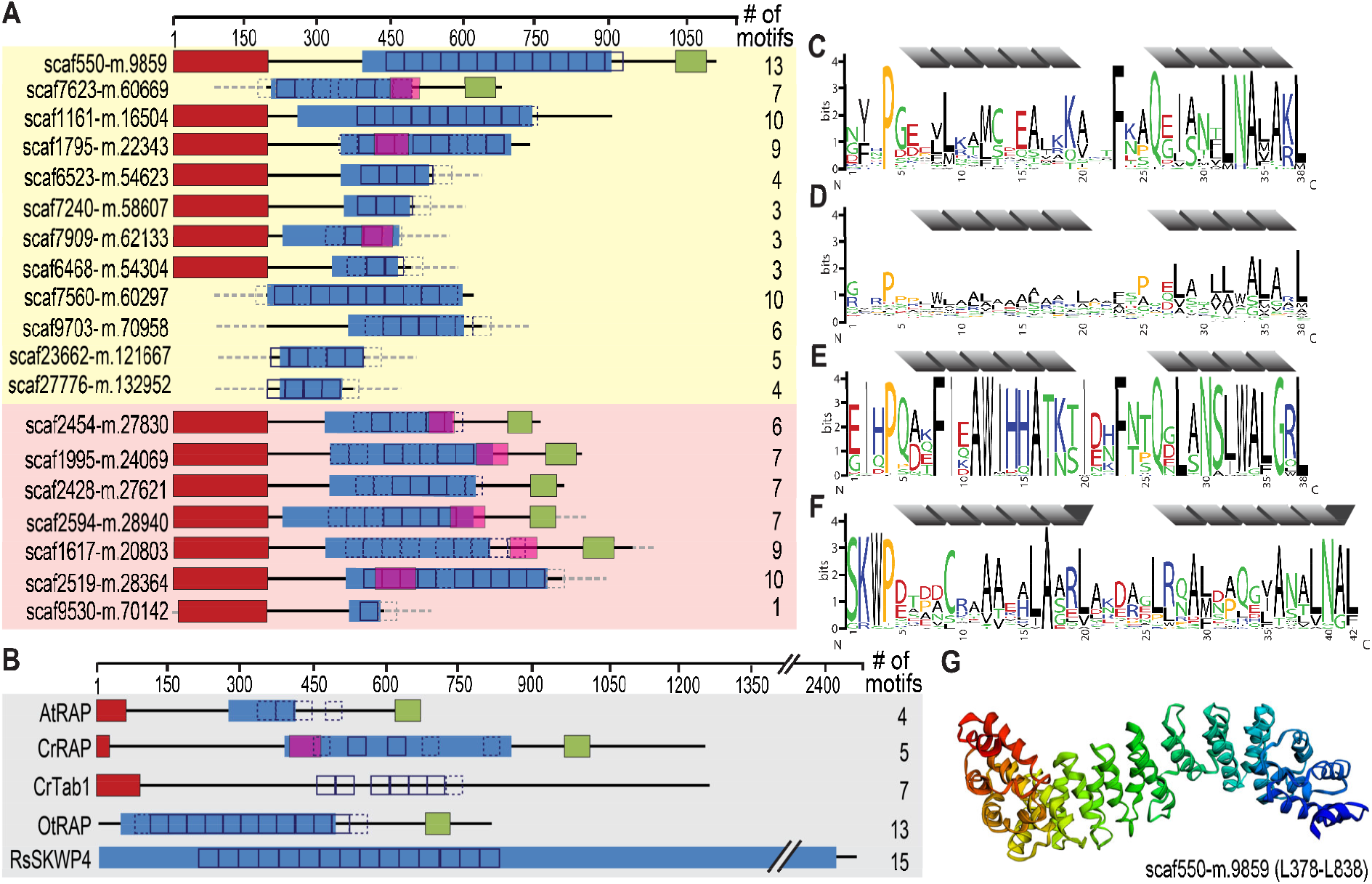
Identification of an expanded family of putative OPR expression regulators targeted to the chromatophore (and mitochondrion) in *P. chromatophora*. **(A)** Domain structure of 12 OPR-containing import candidates identified by MS (yellow background) and further 7 predicted import candidates with a similar domain structure (red background). The number of motif repeats identified in individual proteins is indicated. **(B)** Domain structure and motif repeats in (putative) expression regulators from other organisms. AtRAP, *A. thaliana* RAP domain-containing protein, NP_850176.1, (65); CrTab1, *Chlamydomonas reinhardtii* PsaB expression regulator, ADY68544.1, (92). OtRAP, *Orientia tsutsugamushi* uncharacterized RAP domain-containing protein, KJV97331.1, and RsSKWP4, *Ralstonia soleraceum* RipS4-family effector, AXW63421.1, (93) appear as the highest scoring BlastP/DELTA Blast hits (in the NCBI nr database) for *P. chromatophora* OPR proteins. **(C)** 38-aa repetitive motif found in *P. chromatophora* import candidates. **(D)** OPR motif found in *C. reinhardtii* expression regulators (designed according to (94)). **(E)** Motif derived from *O. tsutsugamushi* OPR proteins. **(F)** 42 aa SKWP motif derived from RipS-family effectors in *R. soleraceum, Xanthomonas euvesicatoria,* and *Mesorhizobium loti* (93, 95, 96). Individual repeats are predicted to fold into two α-helices (grey). Red, targeting signal (crTP for *P. chromatophora* proteins, cTP for AtRAP and CrTab1, mTP for CrRAP); blue, PRK09169-multidomain (Pssm-ID 236394); pink, FAST-kinase like domain (Pssm-ID 310980); green, RAP domain (Pssm-ID 312021); boxes, individual repeats of the motifs shown in C-F (p<e^−20^; p<e^−10^ for CrRAP and CrTab1); dashed boxes, weak motif repeats (p<e^−10^; p<e^−7^ for CrRAP and CrTab1); grey dashed boxes/lines, sequence information incomplete. **(G)** Predicted 3D-structure of the OPR-containing region in scaffold550-m.9859.

Screening the complete *P. chromatophora* transcriptome identified OPR proteins as part of an expanded protein family containing at least 101 members with 1-13 individual OPR motifs (**Table S5**). Besides the 12 chromatophore-localized OPR proteins identified by MS (**Fig. 6A**), of the further 12 OPR proteins identified only in the transcriptome for which full-length N-terminal sequence information was available, seven proteins contained a crTP (**Fig. 6A**), the remaining five a mitochondrial targeting signal.

## Discussion

### Metabolite transport

Despite the obvious need for extensive metabolite exchange between the chromatophore and cytoplasm (12), the chromatophore likely lost on the order of 70 solute transporters following symbiosis establishment (**Fig. 1**). The remaining transport systems do not appear apt to establish metabolic connectivity (**Fig. 2D**). Solely two systems, a DME family and a DASS family transporter, might be involved in metabolite transport. Furthermore, there are three ABC-transporters for which substrate specificity is unknown. However, the high energy costs associated with their ATP-consuming primary active mode of transport appears to be incongruous with high-throughput metabolite shuttling. Some of these ABC-transporters might have become specialized for protein import instead. In line with this idea, the ABC-half transporter PCC0669 that showed highest ion intensities among all chromatophore-encoded transporters (**Fig. 2D**), possesses 33% similarity to *Bradyrhizobium* BclA that functions as an importer for nodule-specific cysteine-rich (NCR) peptides produced by the host plant (38). However, since other transporters in the same family are involved in peroxisomal transport of fatty acids or fatty acyl-CoA (39), similar substrates could also be transported by PCC0669.

In plastids, insertion of nucleus-encoded transporters into the IM is crucial for metabolic connectivity (15–17). Also in more recently established endosymbiotic associations, such as plant sap-feeding insects with nutritional bacterial endosymbionts, multiplication of host transporters followed by their recruitment to the host/endosymbiont interface apparently was involved in establishing metabolic connectivity (40, 41). However, these transporters localize to the symbiosomal membrane, a host membrane that surrounds bacterial endosymbionts. The mechanism enabling metabolite transport across the symbionts’ IM and OM, with symbiont-encoded transport systems being scarce, is a longstanding, unanswered question (42).

Despite the import of hundreds of soluble proteins into the chromatophore, our work provided no evidence for the insertion of nucleus-encoded transporters into the chromatophore IM (or thylakoids). The possibility that such proteins escaped detection for technical reasons appears improbable because: (i) 72% of the chromatophore-encoded transporters were identified in CM samples. Assuming comparable abundances for nucleus-encoded chromatophore-targeted transporters, a large percentage of these proteins should have been detected, too. (ii) More than 100 nucleus-encoded transporters or transporter components were detected in comparable amounts of PM samples showing that our method is feasible to detect this group of proteins. (iii) IM transporters were repeatedly identified in comparable analyses of cyanobacterial (43–47) or plastidial membrane fractions (48–50). Thus, a mechanism to insert nucleus-encoded multi-spanning TM proteins into chromatophore IM and thylakoids likely has not evolved (yet) in *P. chromatophora*.

The protein composition of the chromatophore OM is currently unclear. However, its putative host origin and the notion that proteins traffic into the chromatophore likely via the Golgi (11) suggest that nucleus-encoded transporters can be targeted to the OM by vesicle fusion. Nonetheless, our findings spotlight the puzzling absence of suitable transporters that would allow metabolite exchange across the chromatophore IM. The conservation of active and secondary active IM transporters on the chromatophore genome (**Fig. 2D**) strongly implies that the chromatophore IM kept its barrier function and there is an electrochemical gradient across this lipid bilayer.

In contrast to the absence of multi-spanning TM proteins, we identified numerous short single-spanning TM and AMP-like orphan proteins among chromatophore-targeted proteins. These short import candidates fall into at least four expanded groups, suggesting some degree of functional specialization. Interestingly, expanded arsenals of symbiont-targeted polypeptides convergently evolved in many taxonomically unrelated symbiotic associations and thus seems to represent a powerful strategy to establish host control over bacterial endosymbionts (51). It has been suggested that these ‘symbiotic AMPs’ have the ability to self-translocate across or self-insert into endosymbiont membranes and mediate control over various biological processes in the symbionts including translation, septum formation or modulation of membrane permeability and metabolite exchange (42, 51–56).

The discovery of TMH-containing group 1 proteins appears to be of particular interest in the context of metabolite exchange. The frequent occurrence of (small)xxx(small) motifs might indicate the potential of these proteins to oligomerize (36, 57). The predicted pore-lining residues (**Table S4**) in many of these proteins further suggest that they could form homo- or hetero-oligomeric channels. It has been previously reported that AMPs can arrange in channel-like assemblies which facilitate diffusion along concentration gradients (58, 59), though the lifetime and selectivity of such arrangements requires further investigation. Given the size of the metabolites to be transported, they would be required to form multimer arrangements in barrel-stave (**Fig. S4**), or shortly lived toroidal pores, while maintaining the overall impermeability of the membrane. The formation of such pores still begs the question of how they could maintain a selective metabolite transport. An interesting example in that respect is the VDAC channel of the mitochondrial OM which has been described to follow a stochastic gating mechanism, in which only bigger and, hence, slowly diffusing molecules would be allowed to permeate (60).

An alternative mode of action involves soluble, short import candidates which could interact with the chromatophore envelope membranes via stretches of positively charged aa and amphipathic helices (**Fig. 4A**), and putatively modulate their permeability (42) in what is known as carpet model (61). The mechanism by which such an interaction could cause a transient permeabilization is still a matter of debate, although the asymmetric distribution on the membrane bilayer has been pointed out as plausible reason (62). Further experimental work with the identified proteins could shed light on the potential transport mechanism.

Other short import candidates might also attack intracellular targets. The group 2 sequence motif is found also in hypothetical bacterial proteins which include domains related to DNA processing functions (**Fig. 4B**). Thus, group 2 proteins might provide the host with control over aspects of genetic information processing in the chromatophore. The presence of dozens to hundreds of similar proteins in the various groups, points to a functional interdependence or reciprocal control of individual peptides. In insects, co-occurring AMPs have been shown to synergize, e.g. some AMPs permeabilize membranes to enable entry of other AMPs that have intracellular targets (63).

### Nuclear control over expression of chromatophore-encoded proteins

Besides the establishment of metabolic connectivity, our analyses illuminated another cornerstone in organellogenesis, the evolution of nuclear control over organellar gene expression. Previously, we identified a large number of proteins annotated as transcription factors among chromatophore-targeted proteins (10). Here we described a novel class of chromatophore-targeted helical repeat proteins. Helical repeat proteins appear to represent ubiquitous nuclear factors involved in regulation of organellar gene expression. These proteins are generally characterized by the presence of degenerate 30-40 aa repeat motifs, each of them containing two antiparallel α-helices. The succession of motifs underpins the formation of a super helix that enables sequence specific binding to nucleic acids.

The *P. chromatophora* nuclear genome encodes at least 101 OPR helical repeat proteins (**Fig. 6C**). OPR proteins have mostly been studied in the green alga *C. reinhardtii*, where 44 OPR genes were identified in the nuclear genome. Almost all of these OPR proteins are predicted to localize to organelles [(64); **Fig. 6D**] and five have been shown experimentally to be involved in post-transcriptional steps of chloroplast gene expression. The only known *Arabidopsis* OPR protein is AtRAP [(65); **Fig. 6B**] a factor promoting chloroplast rRNA maturation.

Also the *Paulinella* OPR proteins seem to be mostly organelle-targeted. Many *Paulinella* OPR proteins possess, in addition to the OPR stretches, a Fas-activated serine/threonine (FAST) kinase-like domain (66) and a C-terminal RAP domain (**Fig. 6A**). This domain combination is also present in some of the *Chlamydomonas* OPR proteins (e.g. CrRAP in **Fig. 6B**), the *Arabidopsis* AtRAP protein (**Fig. 6B**), and the FASTK family of vertebrate nucleus-encoded regulators of mitochondrial gene expression (67). Additionally, some bacterial T3SS effector proteins (**Fig. 6B**) show similar domain architectures. However, the exact molecular functions of FAST kinase-like and RAP domains as well as the two conserved regions in *Paulinella* OPR proteins (CR1 and CR2) that share no similarity with known domains remain unknown.

In conclusion, in parallel to the evolution of mitochondria and plastids, also during chromatophore evolution an expanded family of chromatophore-targeted helical repeat proteins evolved. Based on the similarity of their domain architecture to known organelle-targeted expression regulators, the OPR proteins in *P. chromatophora* likely serve as nuclear factors modulating chromatophore gene expression by direct binding to specific target RNAs. Probably chromatophore-targeted OPR proteins evolved from pre-existing mitochondrial expression regulators and were recruited to the chromatophore by crTP acquisition. However, the RNA-binding ability of *Paulinella* OPR proteins, their specific target sequences as well as their ability to modulate expression of chromatophore-encoded proteins remain to be tested experimentally.

## Materials and Methods

### Cultivation of *P. chromatophora* and chromatophore isolation

*P. chromatophora* CCAC0185 [axenic version (7)] was grown (11) and chromatophores isolated as described previously (10). In brief, *P. chromatophora* cells were washed three times with isolation buffer (IB: 50 mM HEPES pH 7.5, 2 mM EGTA, 2 mM MgCl_2_, 250 mM sucrose, 125 mM NaCl) and depleted of dead cells on a discontinuous 20-80% Percoll gradient. The resulting pellet of intact cells was resuspended in IB, cells were broken in a cell disruptor (Constant Systems) at 0.5 kbar, and intact chromatophores isolated on another discontinuous 20-80% Percoll gradient. To increase purity, isolated chromatophores were re-isolated from a fresh Percoll gradient. Recovered chromatophores were washed three times in IB, supplemented with protease inhibitor cocktail (Roche cOmplete), frozen in liquid nitrogen, and stored at −80°C until further use.

### Transmission electron microscopy (TEM)

Isolated chromatophores were fixed in IB containing 1.25% glutaraldehyde for 45 min on ice followed by 30 min post-fixation in 1% OsO_4_ in IB at room temperature. Fixed chromatophores were washed, mixed with 14.5% (w/v) BSA, pelleted, and the pellet fixed with 2.5% glutaraldehyde for 20 min at room temperature. The fixed pellet was dehydrated in rising concentrations of ethanol (from 60% to 100% at −20°C) and then infiltrated with Epon using propylene oxide as a transition solvent. Epon was polymerized at 60°C for 24 h. 70 nm ultrathin sections were prepared and contrasted with uranyl acetate and lead citrate according to (68). A Hitachi H7100 TEM (Hitachi, Tokyo, Japan) with Morada camera (EMSIS GmbH, Münster, Germany) operated at 100 kV was used for TEM analysis. Essentially the same protocol was used for intact *P. chromatophora* cells, however, IB was replaced by growth medium [WARIS-H (69) supplemented with 1.5 mM Na_2_SiO_3_].

### Protein fractionation

#### CM and PM samples

Isolated chromatophores or *P. chromatophora* cells were washed with Buffer I (50 mM HEPES pH 7.5, 125 mM NaCl, 0.5 mM EDTA) at 20,000 ⨯ g or 200 ⨯ g, respectively. Pellets were resuspended in Buffer I and broken by two passages in a cell disrupter at 2.4 kbar. Lysates were supplemented with 500 mM NaCl (final concentration) and passed five times through a 0.6 mm cannula. Cell debris was removed by two successive centrifugations at 15,500 ⨯ g. The supernatant was subjected to ultracentrifugation for 1 h at 150,000 ⨯ g (Beckmann L-80XL optima ultracentrifuge, Rotor 70.1 Ti at 50,000 rpm). Pellets were resuspended in 100 mM Na_2_CO_3_ pH >11 and incubated for 1 h intermitted by 15 passes through a 0.6 mm cannula. Then, insoluble proteins were collected by ultracentrifugation, and subsequently washed with Buffer II (10 mM Tris-HCl pH 7.5, 150 mM NaCl, 0.5 mM EDTA) by passage through a cannula until no particles were visible. Finally, the insoluble fraction was pelleted by ultracentrifugation and solubilized at 36°C in Buffer II supplemented with 1% TritonX-100, 1% Na-deoxycholate, and 0.1% SDS.

#### CL samples

Protein was extracted from intact isolated chromatophores by precipitation with 10% trichloracetic acid for 30 min on ice and pelleted at 21,000 ⨯ g for 20 min. Pellets were washed twice with ice cold acetone for 10 min and finally resuspended in Buffer II plus detergents.

Protein concentration was determined in a Neuhoff assay (70). Aliquots were supplemented with SDS sample buffer (final conc. 35 mM Tris-HCl pH 7.0, 7.5% Glycerol, 3% SDS, 150 mM DTT, Bromophenol blue), frozen in liquid nitrogen, and stored at −80°C until MS-analysis. All steps were performed at 4°C, protease inhibitor cocktail (Roche cOmplete) was added to all buffers used.

### MS analysis and protein identification

Sample preparation and subsequent MS/MS analysis of 3 independent preparations of CM, PM, and CL samples was essentially carried out as described (10). Briefly, proteins were in-gel digested in (per sample) 0.1 μg trypsin in 10 mM ammonium hydrogen carbonate overnight at 37°C and resulting peptides resuspended in 0.1% trifluoroacetic acid. Two independent MS analyses were performed. In MS experiment 1, 500 ng protein per sample, and in MS experiment 2, 500 ng protein per lysate and 1.5 μg protein per membrane sample was analyzed. Peptides were separated on C18 material by LC, injected into a QExactive plus mass spectrometer, and the mass spectrometer was operated as described (10). Raw files were further processed with MaxQuant (MPI for Biochemistry, Planegg, Germany) for protein identification and quantification using standard parameters. MaxQuant 1.6.2.10 was used for the MS experiment 1 analysis and MaxQuant 1.6.3.4 for MS experiment 2. Searches were carried out using 60,108 sequences translated from a *P. chromatophora* transcriptome and the 867 translated genes predicted on the chromatophore genome (10). Peptides and proteins were accepted at a false discovery rate of 1%. Proteomic data have been deposited to the ProteomeXchange Consortium via the PRIDE (71) partner repository with the dataset identifier PXD021087.

### Protein enrichment analysis

Intensities of individual proteins were normalized by division of individual intensities in each replicate by the sum of intensities of all proteins identified with ≥2 peptides in the same replicate. Each protein was assigned an intensity level representing its log10 transformed mean normalized intensity from three replicates in either fraction added 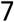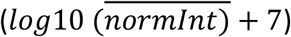, enabling a simple ranking of intensities in a logarithmic range from 0 to 6.

The enrichment factor for each protein in CM as compared to PM or CL samples (*E*_*CM/PM*_ or *E*_*CM/CL*_, respectively) was calculated as 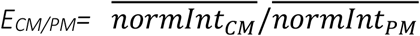 or 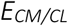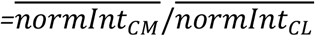 [**Table S2;** missing values (intensity = 0) were excluded from the calculation of means]. Proteins with ≥3 SpC in the chromatophore (i.e. CM+CL fractions) and either *E*_*CM/PM*_>1.5 in at least one of two MS experiments or 0.5<*E*_*CM/PM*_<1.5 in both MS experiments were considered as enriched in chromatophores (see **Fig. S2**). Correspondingly, *E*_*CM/CL*_>1 indicate protein enrichment, *E*_*CM/CL*_<1 depletion in CM samples.

Furthermore, a statistic approach was applied to visualize differences between proteins enriched or exclusively found in a certain fraction. In pairwise comparisons, only proteins were considered showing valid *normInt* values in all three replicates of at least one of the samples being compared. *NormInt* values were log2 transformed and missing values imputed by values from a down shifted normal distribution (width 0.3 SD, down shift 1.8 SD) followed by a pairwise sample comparison based on Student’s t-tests and the significance analysis of microarrays algorithm (S_0_ = 0.8, FDR 5%) (72). Differences between individual proteins in CM vs. PM or CM vs. CL samples were calculated as 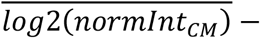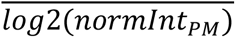 or 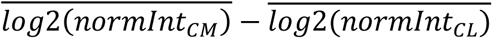, respectively.

### Sequence and structural bioinformatics analyses

TMHs were predicted with TMHMM 2.0 (73), pore-lining residues were predicted with MEMSAT-SVM-pore (74), and AMP peptides were predicted with AmpGram (75). Sequence motifs were discovered using MEME 5.0.5 algorithm (76) in classic mode and visualized with WebLogo (77), number and position of motifs in protein sequences were determined with MAST 5.0.5 using default settings (78). The *P. chromatophora* transcriptome was screened for (i) conserved motifs shown in **Fig. 4A** group 2 and 3 and (ii) the degenerate 38 aa motif shown in **Fig. 6C** using FIMO 5.0.5 with default settings (79). Proteins that contain at least 5 repeats of the 38 aa motif with a p-value < e^−10^ and / or at least 1 repeat with a p-value < e^−20^ were considered candidate OPR-proteins. α-helices in repetitive elements or AMP-like proteins were predicted with Jpred4 (80) and NetSurfP-2.0 (81), respectively. Helical wheel projections were created with HeliQuest (82). Functional protein domains were found with DELTA-BLAST (83). Targeting signals were predicted with PredAlgo (84) for CrRAP and CrTab1, and TargetP 2.0 (85), WoLFPSORT (86), and Predotar (87) for *P. chromatophora* proteins. Tertiary structure predictions were obtained using Phyre2 (88) in normal mode. Area-proportional Venn diagrams were calculated with eulerAPE (89).

Transporters were classified according to the Transporter Classification Database (90). Complete lists of the transporters depicted in Figures 1 and 2D and methods for their identification and classification are provided in Table S1. No OM porins could be identified in the chromatophore genome based on sequence similarity or topology predictions using MCMBB (91).

## Acknowledgments

This study was supported by Deutsche Forschungsgemeinschaft CRC 1208 project B09 (to E.C.M.N.), A03 (to H.G.), and project Z01 (to K.S.). The core facility Elektronenmikroskopie UKD (HHU Düsseldorf) is acknowledged for help with thin sectioning and TEM.

